# Competition among *Aedes aegypti* larvae Kurt Steinwascher

**DOI:** 10.1101/385914

**Authors:** Kurt Steinwascher

## Abstract

Adult *Aedes aegypti* mosquitoes are important vectors of human disease. The size of the adult female affects her success, fitness, and ability to transmit diseases. The size of the adults is determined during the aquatic larval stage. Competition among larvae for food influences the size of the pupa and thus the adult. In these experiments, the food level (mg/larva) and the density (larvae/vial) both affect intraspecific competition, which shows up as the interaction of the two factors. Furthermore, the total food per vial affects the nature of competition among the larvae, also apparent in the interaction of food and density. Male larvae are affected by the percent of males in the vial, but females are not. Seven biologically significant dependent variables were examined, and the data analyzed by multivariate analysis of variance to gain insight into the relationships among the variables and the effects of these factors on the larvae as they grew in small containers. Male and female larvae compete differently from one another for the particulate yeast cells in this experiment; female larvae outcompete males through larger size and by retaining cells within their gut at low total food levels. Under conditions of more intense competition, the pupal masses of both males and females are smaller, so the effect of competition is a reduced apparent food level. The age at pupation is also affected by food and density. Across the twenty treatment combinations of food/larva and larvae/vial, female larvae grew as though there were six different ecological environments while male larvae grew as though there were only four different environments. No interference competition was observed. Eradication efforts aimed at adult populations of this mosquito may inadvertently increase the size and robustness of the next generation of larvae, resulting in a subsequent adult population increase in the second generation.

## Introduction

The *Aedes aegypti* mosquito is a global vector of human diseases, including Yellow Fever, Dengue and Zika. Its impact on human health is through the bite of the adult female; the size and success of the adults are determined by environmental conditions during the larval growth phase ending at pupation [1]. *Aedes aegypti* larvae occur in nature in low numbers spread across multiple small containers [2-13], but see [14].The mosquito larvae react to their environmental conditions including food level (food/larva), total food (food/container), and density (larvae/container) differently depending on gender. Notwithstanding decades of study, including research aimed at developing eradication methods for these and other mosquito species, little is known about the mechanisms by which these larvae interact and compete in the juvenile stages during which they are confined to small containers with limited food resources. Part of the effect of larval density on competition shows up as an apparent change in the food level for mosquitoes [2,6,15-27] and other organisms [28–31]. Investigation of such interactions between food level and density leads to an understanding of the processes underlying the competition among individuals [29–31]. The two sexes of *A. aegypti* respond differently to food level and density [2,4,15–17,19–21,32-39]. Sex differences in the joint effects of food and density suggest that males and females compete for food differently. The experiments in this study explore the response of male and female larvae to different combinations of initial food level, density and the percent of males in each vial. They differ from prior experiments because seven biologically significant dependent variables were measured and analyzed in a single MANOVA, allowing insight into the relationships among the variables as well as the effects of the treatments and most importantly the interactions across the treatments.

## Methods

Eggs were obtained from a colony of *A. aegypti* after feeding females on a mouse. The colony had been started two generations previously with larvae and pupae collected from tires near Dade County Public Works Department (Florida). This research was conducted according to the standard guidelines at the time (1979-1982), sanctioned by the NIH, and under the supervision of the appropriate personnel at the Florida Medical Entomology Laboratory (IFAS and the University of Florida at Gainesville). The food x density experiment investigated the effects of food and density on mosquito larval growth at four different Food levels and five Densities. Numbered, flat-bottomed, shell vials were filled with 20 ml distilled water containing a concentration of baker’s yeast to produce the Food level treatments.

Food levels in the experiment were chosen to span the region where exploitative competition is important: 2 mg, 3 mg, 4 mg, and 5 mg of yeast per larva [17,40–47]. Two hours after eggs were immersed in distilled water, larvae were counted into the numbered shell vials to produce densities of four, five, six, seven, or eight larvae per vial. Five replicates of the twenty density and food level treatments were initiated. Vials were arranged in a randomized sequence, then left in a room at ambient temperatures (18° C to 33° C). Vials were examined for pupae daily from the fourth day through the thirty-seventh day when the last larva died. Pupae were removed from the treatment vial by dropper, blotted on paper toweling, weighed to the nearest 0.01 mg and then identified by sex with a stereo microscope at 10 X magnification.

The sex ratio experiment examined the effect of the percent males and food level on mosquito larval growth to understand competitive interactions between the sexes. Treatment conditions were selected so that survival would be high; vials with less than full survivorship cannot be assigned a sex ratio, nor do they fit a food/larva category, consequently, data from those vials were discarded. Larvae were reared at two densities, five or six larvae per vial, and at two food levels, 3 or 4 mg yeast per larva. Forty replicates of each of the four treatments were initiated. The two densities produced 9 possible mixed-sex ratios at two food levels. The vials were placed in an insectary at 26° C and 12/12 light/dark cycle for the first four days and overnight thereafter. With this exception, handling was identical to the first experiment.

The endpoint of both experiments for each individual larva was either pupation, or death. The endpoint of each treatment vial was the last pupation or the death of the last larva. Seven variables were calculated for each replicated vial of each treatment. These variables were: % survival, mass and age of the Prime male at pupation, Average mass of males at pupation, mass and age of the Prime female at pupation, and Average mass of females at pupation. In each vial, one male and one female were designated as Prime individuals; within that vial each had the greatest expectation of reproductive success for its sex [1]. The Prime individuals, through chance, inherent ability or a combination, appear to be the most successful at larval competition. Because the relationship between pupal mass and age at pupation and adult success differed for the two sexes, the definition of the Prime individual must differ across sexes also. The Prime male was the male with the greatest growth rate (the first male to pupate or the largest of the males pupating on the first day of pupation). For males, an early age at pupation may confer as large an advantage as an increased mass [1]. The Prime female was the largest female to pupate; age is not as important to the success of female, but mass is directly related to fecundity [1]. To compare the growth rates of the Prime individuals, mass and age at pupation were recorded for both sexes. Growth rates are an important indicator of the outcome of exploitative competition among mosquito larvae and other filter feeders [31,48–53], and have been used extensively to score the outcome of intraspecific and interspecific competition among mosquito larvae [1,20,21,26,37,54-63].

Percent survival was calculated to compare the lethal effects of competition among the various treatments and to estimate the relative importance of these lethal effects on the non-lethal changes in mass and age relationships. Data from vials which produced pupae of only one sex were not used in the analysis and this accounts for the variation in sample size among treatments in the food x density experiment. [S1 Table.]

The seven variables were analyzed as a multivariate data set using a two-way analysis of variance design by the program, UNCPROG MANOVA (at the Triangle Universities Computation Center), which accommodates unequal sample sizes. See [64,65]; for a discussion of multivariate analysis of variance.

## Results

### The food x density experiment

The data were analyzed by MANOVA. The individual ANOVA tables were a product of the MANOVA analysis and these were used to further investigate the significant relationships among the 7 dependent variables and across the 9 significant contrasts.

### Multivariate analysis (MANOVA)

The multivariate analysis of the treatment effects and interactions across the seven biologically significant dependent variables should identify the way that competition changes as the initial density and food level vary. For the 9 significant MANOVA contrasts, the correlations between the discriminant function scores and variables [64,65], the R squared values, and the multivariate significance levels are shown in Table 1. The magnitude and sign of the correlations in Table 1 indicate the contribution of each univariate comparison to the significant MANOVA relationship. In the same way that the r squared values guide the selection of the most important contrasts in univariate analyses, the R squared values indicate the relative importance of the multivariate contrasts. There are nine MANOVA contrasts with R squared values greater than 0.60; the remainder are 0.36 or lower. These are distributed across the food level contrasts, the density contrasts and three of the interactions. [S2 Table shows all 19 individual contrasts.]

**Table 1.** Correlations between composite scores and variables with MANOVA significance levels and R squared for the 9 most significant contrasts.

| Contrast (one DF for each) | Survival | Prime male mass at pupation | Prime male age at pupation | Average male mass at pupation | Prime female mass at pupation | Prime female age at pupation | Average female mass at pupation | MANOVA P< | R squared |
| --- | --- | --- | --- | --- | --- | --- | --- | --- | --- |
| **FOOD LEVEL (mg per larva per vial)** |  |  |  |  |  |  |  |  |  |
| **F1: (2 mg + 3 mg) vs (4 mg + 5 mg)** |  | 0.53 |  | 0.61 | 0.77 |  | 0.71 | 0.001 | 0.92 |
| **F2: (2 mg + 4 mg) vs (3 mg + 5 mg)** |  | 0.59 | 0.28 | 0.69 | 0.69 |  | 0.64 | 0.001 | 0.86 |
| **F3: (2 mg + 5 mg) vs (3 mg + 4 mg)** | 0.36 | 0.62 |  | 0.64 | 0.63 |  | 0.53 | 0.001 | 0.84 |
| **DENSITY (larvae per vial)** |  |  |  |  |  |  |  |  |  |
| **D2: 7 larvae vs 8 larvae** |  | 0.70 | 0.43 | 0.67 | 0.39 | 0.30 | 0.39 | 0.001 | 0.68 |
| **D3: (4 + 5 larvae) vs (7 + 8 larvae)** |  | 0.73 | 0.39 | 0.57 | 0.59 | 0.37 | 0.48 | 0.001 | 0.64 |
| **D4: 6 larvae vs (4 + 5 + 7 + 8 larvae)** |  | 0.76 | 0.33 | 0.75 | 0.57 |  | 0.52 | 0.001 | 0.60 |
| **FOOD LEVEL X DENSITY Interactions** |  |  |  |  |  |  |  |  |  |
| **F1 X D3** | 0.25 | 0.63 |  | 0.72 | 0.56 |  | 0.49 | 0.001 | 0.88 |
| **F2 X D1** | 0.28 | 0.55 | 0.25 | 0.62 | 0.62 |  | 0.56 | 0.001 | 0.75 |
| **F2 X D3** | 0.26 | 0.66 |  | 0.75 | 0.53 |  | 0.45 | 0.001 | 0.75 |

There are two patterns apparent from examining Table 1; there are few significant correlations for either Survival, or the Prime female age at pupation, and the correlations that are present are numerically low. Survival and Prime female age at pupation are not affected by the treatments as much as the other dependent variables within the MANOVA.

### MANOVA—food level

Three of the four highest R squared values are associated with the food level contrasts, F1, F2, and F3. It is not surprising that food level is significant, but there are two patterns within these numbers that are striking. First, food level has no correlation with the Prime female age at pupation and almost no correlation with the Prime male age at pupation or Survival. Second, the correlations with the mass variables are all positive, so increased food per larva increases all the mass variables, with notable differences between the two sexes. The Prime female mass has a higher correlation to the food level treatments than the Average female mass, but the Prime male mass has a lower correlation to the food level treatment than the Average male mass. In F1, the contrast with the highest R squared in the table (.92), and the easy-to-understand comparison between the two lower food levels and the two higher food levels, the Prime female mass correlation is 0.77, while the Average female mass correlation is 0.71, and the corresponding correlations for Prime and Average male masses are 0.53 and 0.61. Increased food per larva affects the Prime female more than the Average female, and all females more than the males. However, increased food per larva affects the Average male mass more than it does the Prime male.

### MANOVA—density

Density has no impact on Survival; there are no significant correlations for Survival against any of the density contrasts. Three of the density contrasts, D2, D3, and D4, have high R squared values. D1, the comparison between 4 larvae per vial and 5 larvae per vial, is not significant in the MANOVA, so there is effectively no difference between these two density treatments. For the three other density contrasts, the correlations are all positive; increased density increases the mass variables and to a lesser extent the age at pupation variables. In these contrasts, the Prime male mass at pupation has the largest correlation (0.70, 0.73, 0.76, respectively) with similar scores for the Average male mass. The correlations for the Prime female and Average female masses are lower and in some cases, much lower. Increased density has a larger effect on males than on females (for both mass and age at pupation). According to this multivariate analysis, increases in food level and density both increase the mass at pupation of mosquito larvae. Females are affected by food level more than males and males are affected by density more than females. Increased density increases Age at pupation for males and females, but food level has almost no effect.

### MANOVA—interactions

Interactions between the two main treatments indicate that the observed results are higher or lower than would be expected based on the effects of the main treatments. The interaction with the highest R squared value (0.88) is F1 X D3, where F1 is the contrast between the two lowest food levels against the two highest food levels, and D3 is the contrast between the two lowest densities against the two highest densities. This interaction does have a correlation with survival, although it is small (0.25). The interaction does not have a correlation with either the Prime male age at pupation or the Prime female age at pupation. The four mass variables show large positive correlations with this interaction; the Average male mass has the highest (0.72) followed by the Prime male mass (0.63), the Prime female mass (0.56) and the Average female mass (0.49). This interaction affects males more than females, and the Average male mass more than the Prime male mass, but Prime female mass more than Average female mass. The four cells of this contrast represent the most extreme competition (the two lowest food levels with the two highest densities), the least extreme competition (the two highest food levels with the two lowest densities), the highest total food per vial (the two highest food levels with the two highest densities) and the lowest total food per vial (the two lowest food levels with the two lowest densities). The significance of this interaction indicates that either competition, or total food per vial, or both influence the growth of the larvae in the microcosms.

The other two interactions with high R squared values (both 0.75) are F2 X D1 and F2 X D3, where F2 is [(2 mg/larva + 4 mg/larva) vs (3 mg/larva + 5 mg/larva)], D1 is the contrast between 4 larvae and 5 larvae, and D3 is the contrast between the two lowest densities and the two highest densities, as before. Both interactions have small correlations with survival (0.28 and 0.26). Neither interaction has a correlation with Prime female age at pupation and only F2 X D1 has a small correlation with Prime male age at pupation. However, both interactions have similar significant correlation with the four mass variables as the F1 X D3 interaction above. For F2 X D1, the correlation for the Average male mass (0.62) is greater than the Prime male mass (0.55) and that of the Prime female mass (0.62) is greater than that of the Average female mass (0.56). For F2 X D3, the correlation for the Average male mass (0.75) is also greater than the Prime male mass (0.66) and that of the Prime female mass (0.53) is similarly greater than that of the Average female mass (0.45).

According to the multivariate interactions, males (mass variables) are affected more by the interactions than females and the Average male is affected more than the Prime male. However, the Prime female is affected more by the interactions that the Average female. In the food x density experiment, the MANOVA correlations for the main food level contrasts show one pattern, the correlations for the main density contrasts show a separate pattern, and the three significant interactions show a third pattern.

To understand what these significant interactions mean to the biology of *Aedes aegypti*, we need to examine the contrasts in the univariate analyses.

### MANOVA summary

There are four different patterns for the seven dependent variables across the treatment contrasts. All the mass variables have significant positive correlations with the main treatments and with the three significant interactions. Food level and density both affect the mass of the larvae, but the significant interactions suggest that the effect of density may be through differences in the amount of food available (total food per vial) or through competitive interactions or both, so we can’t tell the magnitude of independent effects of food level and density on the growth of the larvae.

The second pattern is for the Prime female age at pupation. The only significant contrasts for which this variable had a correlation in the MANOVA were two associated with density. For Prime female age at pupation, there is an independent effect of density; higher density increases the age at pupation for the Prime female, but none of the food level treatments or interactions affect it.

The third pattern is for the Prime male age at pupation. It is affected by food level, density and the interactions, but it appears to be much more strongly affected by density than by either food level or the interactions. This suggests that there is an independent effect of density on the Prime male age at pupation, and a smaller effect of food level that interacts with density at the lowest densities (D1: 4 larvae vs 5 larvae). Different factors determine the male age at pupation than those that determine the female age at pupation.

The fourth pattern is for Survival. This variable is affected by food level and by the interactions, but not by the main density treatment. This suggests that part of the effect of food level on Survival is mediated by the total food per vial.

### Univariate analyses—ANOVAs

Table 2 presents the r squared values for each dependent variable summed across each of the main treatments and all the interactions. The last row of Table 2 shows the total r squared for each of the seven variables. These range from 0.46 (Prime female age at pupation) to 0.94 (both Average male mass and Average female mass). What is noteworthy here is that the variation in Prime female age at pupation is not well explained by the food level and density treatments; only 46 % of the variation in Prime female age at pupation is explained by the treatments, 54 % is due to factors not included in the experiment. More interesting is that the variation in the four mass variables is very well explained by the experimental treatments (88 %, 94 %, 93 %, and 94 %). Prime male age at pupation and Survival are in between these extremes (69 % and 63 %, respectively). [S3 Table shows the Mean Square value, the significance (P value), and the r squared value for each of the 19 contrasts and each of the seven dependent variables. This is sufficient to construct the individual ANOVAs for the dependent variables.]

**Table 2.** r squared values summed across treatments and interactions for each of the 7 dependent variables.

| Treatments | DF | Survival | Prime male mass at pupation | Prime male age at pupation | Average male mass at pupation | Prime female mass at pupation | Prime female age at pupation | Average female mass at pupation |
| --- | --- | --- | --- | --- | --- | --- | --- | --- |
| Food Level | 3 | 0.30 | 0.41 | 0.21 | 0.47 | 0.64 | 0.12 | 0.64 |
| Density | 4 | 0.00 | 0.16 | 0.21 | 0.12 | 0.06 | 0.20 | 0.08 |
| Food Level X Density Interactions | 12 | 0.33 | 0.31 | 0.27 | 0.35 | 0.23 | 0.14 | 0.22 |
| Totals | 19 | 0.63 | 0.88 | 0.69 | 0.94 | 0.93 | 0.46 | 0.94 |
| Treatments | DF | Survival | Prime male mass at pupation | Prime male age at pupation | Average male mass at pupation | Prime female mass at pupation | Prime female age at pupation | Average female mass at pupation |
| Food Level | 3 | 0.30 | 0.41 | 0.21 | 0.47 | 0.64 | 0.12 | 0.64 |
| Density | 4 | 0.00 | 0.16 | 0.21 | 0.12 | 0.06 | 0.20 | 0.08 |
| Food Level X Density Interactions | 12 | 0.33 | 0.31 | 0.27 | 0.35 | 0.23 | 0.14 | 0.22 |
| Totals | 19 | 0.63 | 0.88 | 0.69 | 0.94 | 0.93 | 0.46 | 0.94 |

### ANOVAs—food level

The treatments food level and density are expected to have significant effects on these variables based on prior experiments. The treatment conditions were selected to produce different levels of non-lethal competition. The MANOVA indicated that there were differences in the way that the dependent variables responded to the treatments and interactions. We see in Table 2, that food level alone accounts for 30 % of the variation in Survival, more than 40 % of the variation in the two male mass variables and 64 % of the variation in the two female mass variables. Females (mass variables) are much more affected by food level than are males. While food level explains the same amount of variation in the two female mass variables (64 %), it accounts for more of the variation in Average male mass (47 %) than in the Prime male mass (41 %). In contrast to the large effect on the female mass variables and the male mass variables, food level accounts for only 12 % of the variation in Prime female age at pupation, and 21 % of Prime male age at pupation.

Survival is higher at the intermediate food levels (3 mg/larva, 4 mg/larva) than at the highest (5 mg/larva) or lowest (2 mg/larva) [S4 Table]. This is likely the reason for the low correlation for Survival against food level in the MANOVA. [Contrast F3: (2 mg/larva + 5 mg/larva) vs (3 mg/ larva + 4 mg/larva) is the only significant correlation for Survival against food level in the MANOVA.]

The Prime female mass at pupation, the Average female mass at pupation, the Prime male mass at pupation and the Average male mass at pupation all increase with increasing food/larva [S5 Table, S6 Table, S7 Table, S8 Table]. The effect of Food level on these mass variables is consistent with the relationships described by the correlations in the MANOVA.

The Prime male age at pupation is highest at the highest food/larva treatment; the other food level treatments are lower, but similar to each other [S9 Table]. The Prime female age at pupation is highest at the highest food/larva treatment and lowest at the next highest food/larva treatment (4 mg/larva) [S10 Table]. Neither of the age at pupation variables had large correlations with food level in the MANOVA.

### ANOVAs—density

The next row in Table 2 shows the contribution of the density treatments to the total r squared values. The highest r squared values are for the Prime male age at pupation (21 %) and the Prime female age at pupation (20 %). Density affects the male mass variables (16 %, 12 %) more than the female mass variables (6 %, 8 %); and affects the sexes differently. Density explains more of the variation in the Prime male mass than in the Average male mass, but explains more of the variation in the Average female mass than in the Prime female mass.

Survival is unaffected by density (r squared = 0.00). This is consistent with the zero correlation with density observed in the MANOVA result.

The Prime male age at pupation is highest at the highest density; the lower densities are similar in age at pupation [S9 Table]. The Prime female age at pupation is also highest at the highest density; the lower densities vary, but with no obvious pattern [S10 Table]. This may be the (lack of) pattern that resulted in the low positive correlations for age at pupation with density in the MANOVA.

Prime male mass at pupation is highest at the highest Density and similar at lower densities [S7 Table]. Average male mass at pupation is highest at the lowest Density and similar at higher densities [S8 Table]. Prime female mass at pupation increases from the lowest density to the second highest density, but then is lowest at the highest density [S5 Table]. Average female mass at pupation is lowest at the highest density, and the middle density (6 larvae/vial), but similar in the other density treatments [S6 Table]. There is no uniform effect of density across the different mass variables. All the mass variables had significant interactions between food level and density in the MANOVA, so the effect of density on the mass variables may be mediated by total food per vial, competition or both.

The ANOVA reveals the effect of treatments on the individual variables, while the MANOVA reveals the relationships among the variables for each of the contrasts. The MANOVA indicated that the density treatments affected the male mass variables more than the female mass variables. The ANOVA reflected that result as well.

The MANOVA indicated that density affected Prime male mass more than Prime male age at pupation, but the ANOVA explained the Prime male age at pupation more than the Prime male mass. The observed result for females is similar. There is a component of the age at pupation in the ANOVA that is independent of the Density treatment correlations calculated by the MANOVA.

### ANOVAs—interactions

The third row in Table 2 shows the contribution of the food level X density interactions to the r squared values. More than half of the contribution to the Survival r squared total is due to the interactions (33 %). Interactions explain more than 30 % of the variation in the male mass variables and more than 20 % of the variation in the female mass variables. Interactions explain more of the variation in Prime male age at pupation (27 %) than in Prime female age at pupation (14 %).

In Table 3 the single degree of freedom interaction contrasts with the largest r squared values are the same contrasts that were identified in the MANOVA: F1 X D3, F2 X D1, and F2 X D3. These three contrasts account for most of the r squared value in the overall Food level X Density interactions (Table 3, last row, Totals, compared to Table 2, row labelled Food level X Density Interactions). We know that both food level and density affect six of the seven dependent variables in this experiment (Table 2). The interaction between food level and density is significant when the Mean Squares are higher or lower than expected due to the main effects of food level and density separately. These patterns should help us understand the biological interactions among the larvae in the vials. As mentioned before, these three contrasts show the effects of competition and total food per vial on the growth of the larvae in the microcosms.

**Table 3.** r squared values summed across the three main interactions for each of the 7 dependent variables.

| Food Level X Density<br>Interaction contrasts<br>(single DF) | DF | Survival | Prime male<br>mass at<br>pupation | Prime male<br>age at<br>pupation | Average<br>male mass<br>at pupation | Prime female<br>mass at<br>pupation | Prime female<br>age at<br>pupation | Average<br>female mass<br>at pupation |
| --- | --- | --- | --- | --- | --- | --- | --- | --- |
| **F1 X D3** | 1 | 0.13 | 0.15 | 0.08 | 0.17 | 0.11 | 0.09 | 0.10 |
| **F2 X D1** | 1 | 0.07 | 0.05 | 0.04 | 0.05 | 0.06 |  | 0.06 |
| **F2 X D3** | 1 | 0.06 | 0.07 | 0.03 | 0.09 | 0.05 |  | 0.04 |
| **Totals** | 3 | 0.26 | 0.27 | 0.15 | 0.31 | 0.22 | 0.09 | 0.20 |

Tables 4-7 present the means and standard errors for the three interaction contrasts: F1 X D3, F2 X D1, and F2 X D3. F1 compares the low food levels (2 mg/larva + 3 mg/larva) with the high food levels (4 mg/larva + 5 mg/larva) and D3 compares the low densities (4 larvae/vial + 5 larvae/vial) with the high densities (7 larvae/vial + 8 larvae/vial). F2 compares the low food levels (2 mg/larva + 4 mg/larva) with the high food levels (3 mg/larva + 5 mg/larva) and D1 compares 4 larvae/vial with 5 larvae/vial. The vials with the low densities and high food levels should experience the least competition and the vials with the high densities and low food levels should experience the most competition. The other treatments should experience levels of competition in between the extremes, and represent the vials with the least and most total food per vial. The value of Average male mass at pupation is highest at the low density: high food levels treatment (the least competition) and lowest at the high density: low food levels combination (the most competition) and intermediate at the other combinations (intermediate levels of competition). However, none of the other variables show this pattern.

**Table 4.** Comparison of Means and (Standard Errors) for the 3 significant interactions for Survival.

| Treatment |  | Survival<br>F1 X D3 | Survival<br>F2 X D1 | Survival<br>F2 X D3 |
| --- | --- | --- | --- | --- |
| **Least competition** | Low Density—D1=(4 larvae); D3=(4 larvae + 5 larvae)<br>High Food Level— F1=(4 mg/larva + 5 mg/larva);<br>F2=(3 mg/larva + 5 mg/larva) | 1.25 (0.26) | 1.32 (0.16) | 1.15<br>(0.24) |
| **Least food/vial** | Low Density—D1=(4 larvae); D3=(4 larvae + 5 larvae)<br>Low Food Level— F1=(2 mg/larva + 3 mg/larva);<br>F2=(2 mg/larva + 4 mg/larva) | 0.91 (0.31) | 0.95 (0.61) | 1.01<br>(0.41) |
| **Most food/vial** | High Density— D1=(5 larvae); D3=(7 larvae + 8 larvae)<br>High Food Level— F1=(4 mg/larva + 5 mg/larva);<br>F2=(3 mg/larva + 5 mg/larva) | 1.05 (0.35) | 0.98 (0.16) | 1.00<br>(0.33) |
| **Most competition** | High Density— D1=(5 larvae); D3=(7 larvae + 8 larvae)<br>Low Food Level— F1=(2 mg/larva + 3 mg/larva);<br>F2=(2 mg/larva + 4 mg/larva) | 1.03 (0.29) | 1.07 (0.34) | 1.08<br>(0.30) |
| **r squared value;**<br>**\*\*\* = P<0.001** |  | 0.13*** | 0.07*** | 0.06*** |

**Table 5.** Comparison of Means and (Standard Errors) for the 3 significant interactions for the Age at Pupation variables.

| Treatment |  | Prime male<br>age at<br>Pupation<br>F1 X D3 | Prime male<br>age at<br>Pupation<br>F2 X D1 | Prime male<br>age at<br>Pupation<br>F2 X D3 | Prime<br>female age<br>at pupation<br>F1 X D3 | Prime<br>female age<br>at pupation<br>F2 X D1 | Prime<br>female age<br>at pupation<br>F2 X D3 |
| --- | --- | --- | --- | --- | --- | --- | --- |
| **Least competition** | Low Density—D1=(4 larvae); D3=(4 larvae + 5 larvae)<br>High Food Level— F1=(4 mg/larva + 5 mg/larva); F2=(3 mg/larva + 5 mg/larva) | 5.59 (0.66) | 5.55 (0.07) | 5.73 (0.53) | 6.90 (0.96) | 7.60 (0.28) | 7.88 (0.38) |
| **Least food/vial** | Low Density—D1=(4 larvae); D3=(4 larvae + 5 larvae)<br>Low Food Level— F1=(2 mg/larva + 3 mg/larva); F2=(2 mg/larva + 4 mg/larva) | 5.48 (0.17) | 5.40 (0.14) | 5.38 (0.30) | 7.75 (0.54) | 7.00 (1.41) | 6.80 (0.91) |
| **Most food/vial** | High Density— D1=(5 larvae); D3=(7 larvae + 8 larvae)<br>High Food Level— F1=(4 mg/larva + 5 mg/larva); F2=(3 mg/larva + 5 mg/larva) | 6.32 (1.54) | 5.90 (0.85) | 6.25 (1.61) | 8.43 (2.87) | 8.15 (0.21) | 9.68 (1.80) |
| **Most competition** | High Density— D1=(5 larvae); D3=(7 larvae + 8 larvae)<br>Low Food Level— F1=(2 mg/larva + 3 mg/larva); F2=(2 mg/larva + 4 mg/larva) | 5.50 (0.10) | 5.35 (0.49) | 5.13 (0.15) | 8.38 (1.60) | 6.60 (0.57) | 7.13 (1.80) |
| **r squared value;**<br>**\*\*\* = P<0.001,**<br>**\*\*=P<0.01** |  | 0.08*** | 0.04*** | 0.03** | 0.09*** | ns | ns |

**Table 6.** Comparison of Means and (Standard Errors) for the 3 significant interactions for Male Mass at Pupation.

| Treatment |  | Prime male<br>mass at<br>pupation<br>F1 X D3 | Prime male<br>mass at<br>pupation<br>F2 X D1 | Prime male<br>mass at<br>pupation<br>F2 X D3 | Average<br>male mass<br>at pupation<br>F1 X D3 | Average<br>male mass<br>at pupation<br>F2 X D1 | Average<br>male mass<br>at pupation<br>F2 X D3 |
| --- | --- | --- | --- | --- | --- | --- | --- |
| **Least competition** | Low Density—D1=(4 larvae); D3=(4 larvae + 5 larvae)<br>High Food Level— F1=(4 mg/larva + 5 mg/larva); F2=(3 mg/larva + 5 mg/larva) | 2.39 (0.15) | 2.34 (0.17) | 2.37 (0.17) | 2.42 (0.20) | 2.41 (0.28) | 2.39 (0.23) |
| **Least food/ vial** | Low Density—D1=(4 larvae); D3=(4 larvae + 5 larvae)<br>Low Food Level— F1=(2 mg/larva + 3 mg/larva); F2=(2 mg/larva + 4 mg/larva) | 2.08 (0.21) | 2.14 (0.09) | 2.10 (0.23) | 2.04 (0.23) | 2.13 (0.08) | 2.07 (0.26) |
| **Most food/vial** | High Density— D1=(5 larvae); D3=(7 larvae + 8 larvae)<br>High Food Level— F1=(4 mg/larva + 5 mg/larva); F2=(3 mg/larva + 5 mg/larva) | 2.49 (0.20) | 2.39 (0.23) | 2.43 (0.24) | 2.38 (0.17) | 2.37 (0.28) | 2.32 (0.23) |
| **Most competition** | High Density— D1=(5 larvae); D3=(7 larvae + 8 larvae)<br>Low Food Level— F1=(2 mg/larva + 3 mg/larva); F2=(2 mg/larva + 4 mg/larva) | 2.05 (0.22) | 2.07 (0.39) | 2.11 (0.30) | 1.95 (0.21) | 2.01 (0.43) | 2.01 (0.28) |
| **r squared value;**<br>**\*\*\* = P<0.001** |  | 0.15*** | 0.05*** | 0.07*** | 0.17*** | 0.05*** | 0.09*** |

**Table 7.** Comparison of Means and (Standard Errors) for the 3 significant interactions for Female Mass at Pupation.

| Treatment |  | Prime female mass at pupation<br>F1 X D3 | Prime female mass at pupation<br>F2 X D1 | Prime female mass at pupation<br>F2 X D3 | Average female mass at pupation<br>F1 X D3 | Average female mass at pupation<br>F2 X D1 | Average female mass at pupation<br>F2 X D3 |
| --- | --- | --- | --- | --- | --- | --- | --- |
| **Least competition** | Low Density—D1=(4 larvae); D3=(4 larvae + 5 larvae)<br>High Food Level— F1=(4 mg/larva + 5 mg/larva); F2=(3 mg/larva + 5 mg/larva) | 3.95 (0.32) | 3.72 (0.74) | 3.66 (0.65) | 3.75 (0.31) | 3.53 (0.82) | 3.48 (0.63) |
| **Least food/vial** | Low Density—D1=(4 larvae); D3=(4 larvae + 5 larvae)<br>Low Food Level— F1=(2 mg/larva + 3 mg/larva); F2=(2 mg/larva + 4 mg/larva) | 2.80 (0.35) | 3.04 (0.76) | 3.10 (0.68) | 2.72 (0.25) | 2.98 (0.67) | 3.00 (0.57) |
| **Most food/vial** | High Density— D1=(5 larvae); D3=(7 larvae + 8 larvae)<br>High Food Level— F1=(4 mg/larva + 5 mg/larva); F2=(3 mg/larva + 5 mg/larva) | 4.03 (0.30) | 3.59 (0.85) | 3.77 (0.61) | 3.85 (0.41) | 3.43 (0.70) | 3.58 (0.72) |
| **Most competition** | High Density— D1=(5 larvae); D3=(7 larvae + 8 larvae)<br>Low Food Level— F1=(2 mg/larva + 3 mg/larva); F2=(2 mg/larva + 4 mg/larva) | 2.75 (0.59) | 3.16 (0.91) | 3.01 (0.89) | 2.55 (0.48) | 3.03 (0.71) | 2.83 (0.78) |
| **r squared value;**<br>**\*\*\* = P<0.001** |  | 0.11*** | 0.06*** | 0.05*** | 0.10*** | 0.06*** | 0.04*** |

Another way to rank these treatments is by total food per vial (calculated by multiplying the number of larvae per vial by the food per larvae). None of the variables line up strictly according to total food per vial, but Prime male mass at pupation, Prime male age at pupation, Prime female mass at pupation, Prime female age at pupation, and Average female mass all reach their largest value in the vials with the most food per vial.

Survival is affected by the food level treatments and the interactions, but not at all by the Density treatments (Table 2). In the interaction contrasts (Table 4), Survival is highest at the low density: high food level combinations (least competition) and lowest at the low density: low food level combinations (least total food per vial). The Survival values in the high density treatments are intermediate, but the survival is higher at the higher food level (most total food per vial). Survival is lowest at the lowest total food per vial and increases as the total food increases. This doesn’t entirely explain the variation in Survival because the highest total food per vial is associated with a lower percent survival than the next highest (the treatments with the least competition). This suggests that Survival is affected by both total food per vial and competition among the larvae. In other words, at least part of the effect of density is due to the increase in total food per vial. This doesn’t rule out a separate effect of density independent of food level in the interaction contrasts.

Table 5 compares the two Age at pupation variables. The Prime male age at pupation is very similar across three of the four treatments (5.13 - 5.59 days), but it is much longer (5.90-6.32 days) in the treatments with the highest total food per vial. The Prime male age at pupation does not seem to be affected by competition; the vials with the least competition and those with the most competition are similar, but the vials with intermediate levels of competition and the most food per vial take the longest to pupate. This is not the same pattern for the Prime female age at pupation. First, only the F1 X D3 interaction contrast is significant for this variable. Second, the Prime female takes longer than the Prime male to pupate in all treatments. Third, the Prime female pupates earliest in the treatments with the least competition (6.90 days). Reducing the food level at the lower densities results in later pupation, but increasing the density increases the age at pupation even further (with little difference between the food/larva levels). Clearly males and females are responding to different external or internal conditions to trigger pupation. Males pupate at about 5 1/2 days except when there is a lot of food in the vials; females pupate earliest in the vials with the least competition but seem to be affected by both the density and the food level in the other treatments. The MANOVA indicated that the two Age at pupation variables were not correlated with the four mass variables, so these significant interactions are independent of the behavior of the mass variables.

The four mass variables are examined three ways: 1) individually; 2) within sexes to compare the Prime individual with the Average; and 3) across sexes to compare the two Primes and the two Averages. The MANOVA showed that all four variables had significant positive correlations for each of the three interaction contrasts.

Table 6 presents the means and standard errors for the three significant interactions for both Prime male mass at pupation and Average male mass at pupation. For the Prime male mass at pupation the highest mass values are at the high food levels and the lowest are at the low food levels. The lowest mass value is in the vials with the most competition, but the highest mass value is in the vials with the most food per vial, not the ones with the least competition. For the Average male mass, the lowest value is also in the vials with the most competition, but the highest mass value is in the vials with the least competition. Competition appears to be the main determinant of growth for the Average male. The Prime male was defined as the largest of the first males to pupate in each vial. The mean values of the Prime male mass are larger than the mean values of the Average male mass in all treatments except for the ones with the least competition (low density: high food levels); in these treatments, the Average mass of males is greater than the Prime male mass. This means that the Prime male pupates while there is still enough food for the remainder of the males to continue to grow and pupate at a larger size than the Prime male. Another comparison between the Prime male mass and the Average male mass is the difference between the two values at low density and low food level (0.03 mg - 0.04 mg) compared to the two high density treatments (0.10 mg - 0.11 mg) (F1 X D3 and F2 X D3, Table 6). The distribution of sizes among males is tighter at the lowest total food per vial A greater difference between the size of the Prime male and the Average males at low resource levels would be an indicator of interference competition, thus no interference competition among males is evident here.

For the Prime female mass at pupation the two highest mass values are also at the high food levels and the two lowest are at the low food levels (Table 7). For two of the contrasts the lowest mass value is in the vials with the most competition, but the highest mass value is in the vials with the most food per vial, not the ones with the least competition (F1 X D3 and F2 X D3, Table 7). Unlike the pattern for the Average male mass, the Average female mass mirrors the Prime female mass exactly. The interaction for both the Prime and Average female mass variables is the same as that for the Prime male (above). The Prime female is defined as the largest female to pupate so it is always larger than the Average. Comparing the values of the means of the Prime female mass and the Average female mass, the Prime female is about 0.18 mg - 0.20 mg larger than the average female except in the treatment with the low density and low food levels (0.08 mg - 0.10 mg). These vials have the lowest levels of total food per vial in the experiment. The relative sizes of the Prime females and the Average females are similar across treatments except at the lowest total food per vial, when the relative size difference of the two is much smaller. Again, an increase in the distribution of sizes at low resource levels would be an indicator of interference competition, thus no interference competition among females is apparent in these two contrasts. The distribution of sizes among females is affected by total food per vial, or competition, or both, rather than either food level or density independently.

For the F2 X D1 contrast, the pattern is different from the other two interaction contrasts for both the Prime female and the Average female masses. Prime female mass at pupation is greatest in the vials with the least competition (3.72 mg). The lowest value is in the vials with the least total food (3.04 mg). The Average female mass at pupation shows the same pattern as the Prime female for this contrast. The total food in all the vials in this contrast is at the low end of the total food per vial across the entire experiment. This contrast compares the two lowest densities (4 larvae/vial vs 5 larvae/vial), so the highest total food per vial is going to be 25 mg/vial rather than 40 mg/vial (at the 8 larvae/vial density and 5 mg food per larva). Both density and total food per vial are at the low end of the range of the entire experiment. Within this subset of the experiment, the females in the vial with the least competition grow larger than the females with the most total food per vial, so competition appears to be more important at lower food levels and/or lower levels of total food per vial. Furthermore, the females in the vials with the least total food are smaller than those in the vials with the most competition. The greater total food in the vials with the most competition allows those females to grow larger than in the vials with the least total food, despite the same food/larva in both sets of vials. The least total food per vial, which results in the smallest mass at pupation for both the Prime and Average females, also results in the smallest difference between the Prime and Average females (0.06 mg compared to 0.13 - 0.19 mg for the other treatments). This is similar to the result for the other two interaction contrasts. An increase in the distribution of sizes at low resource levels would be an indicator of interference competition, thus no interference competition among females is apparent even at the lowest food levels. The distribution of sizes among females is affected by total food per vial rather than either food level or density independently.

Summarizing the differences within each sex, the interactions reveal differences between the Prime male mass and the Average male mass with the Prime male growing largest at the highest total food per vial and smallest in the vials with the most competition. The Average male mass is largest in the vials with the least competition and smallest in the vials with the most competition (Table 6). For females, two of the interactions mirror the pattern of the Prime male mass for both Prime female mass and Average female mass. The remaining interaction (F2 X D1) suggests that lower total food per vial affects the competition among females in these vials—a subset of the entire experiment (Table 7).

Comparing the two sexes at pupation (Tables 6 and 7), the Prime female mass is always greater than the Prime male mass, but the difference is larger at high food levels (1.20 mg -1.56 mg) than at low food levels (0.70 mg - 1.09 mg). The Average female mass is similarly greater that the Average male mass, and the difference is also larger at high food levels (1.06 mg - 1.47 mg) than at low food levels (0.60 mg - 01.02 mg), but the difference between the two Averages is always smaller than the difference between the two Prime masses. This indicates that females outcompete males for food within the limits of the food x density experiment.

For the two interactions F1 X D3 and F2 X D3, the difference between the female and male masses is smallest in vials with the most competition followed by the vials with the least total food. The difference in size of the Prime male and female is similar in the vials with the least competition and most total food, but the difference between the Averages is greater for the most total food than for the least competition. For the interaction F2 X D1, the smallest difference between the mass of males and females is in the vials with the least food and the largest difference is in the vials with the least competition. An increase in the distribution of sizes at low resource levels would be an indicator of interference competition, thus no interference competition between males and females is apparent.

### Effect of sex ratio and food on mosquito larval growth

In the sex ratio experiment, the food per larva and the larvae per vial were chosen from the middle of the values for the first experiment: 3 mg/larva or 4 mg/larva, and 5 larvae/vial or 6 larvae/vial. The sex ratio was calculated for each vial with 100 % pupation (100 % survival). As mentioned earlier, neither the sex ratio nor the Food level can be determined accurately if there is any mortality.

For the subset of vials with 100 % pupation, the overall sex ratio was 52 % male. As before, the mass was measured for each pupa and values were calculated for Prime male mass, Average male mass, Prime female mass and Average female mass. Means, standard deviations and sample sizes for the mass of the Prime male are presented in S11 Table arranged by sex ratio (% males) and by food level. The food level obviously affects the size of the Prime male. At both food levels the mass of the Prime male appears to increase with an increase in the % males in the vial. The regression of mass on sex ratio was significant at the lower food level (F(1,18) = 7.869, P < 0.05, r squared = .20), but not at the higher food level (F(1,12) = 3.256, NS). At the lower food level, the mass of the Prime male increases as the percent of males increases. None of the other three mass variables had a significant regression on sex ratio at either food level.

Because males pupate earlier and at a smaller mass than females, an increase in percent of males is expected to correspond to a relative increase in food level. At the lower food level, the mass of the Prime male increases as the percent of males increases; an increase in the percent of males acts as though the food level increased for the Prime male. Since none of the other mass variables responds to sex ratio at either food level, this suggests that the Prime male outcompetes the other males for food. The Prime male mass is included in the Average male mass, so a systematic increase in the Prime male mass with no effect on the Average implies that the non-Prime males are symmetrically decreasing in size. It also suggests that females are unaffected by competition with males, which implies that the two sexes are using the food resource differently.

## Discussion

Mosquito larvae filter particles indiscriminately [48,50] . They can filter particles from the water column, from submerged surfaces (leaves, container walls), and abrade solids (such as dead larvae or other carcasses) into ingestible particles [38,66–68]. Besides discrete particles such as the yeast cells used in these experiments, in nature mosquito larvae may ingest gels, colloids and dissolved nutrients that contribute to their nourishment [66]. In contrast to the indiscriminate filtering, mosquito larvae actively seek out and aggregate at food rich locations, respond to organic chemicals that leach from potential food, change their feeding rate (the beating rate of the lateral palatal brushes), and alter the proportion of time spent feeding in response to hunger, food availability, and neurochemicals [38,67,68]. Many filter feeders, including some mosquito larvae, pass much of the ingested food through the gut intact [48,52,66,69]; only the most available subset of nutrients is assimilated. Normal feeding for *A. aegypti* larvae results in passage through the gut in 0.5-1.0 hour [66]. When food level (mg/larva) decreases, the proportion of the total nutrients assimilated and hence the efficiency (mg larval growth/mg food ingested) can be increased by decreasing the feeding rate and retaining food within the gut for longer intervals in other filter feeders [31,52]. In a pattern analogous to tadpoles, mosquito larvae retain food when transferred into distilled water [48] suggesting that they may vary their feeding rates and efficiencies in response to the availability of food similarly.

Other experiments show that male and female 4th instar larvae are competent to pupate at 24-36 hours after they molt from the 3rd instar [70–73], but that high food levels cause them to delay pupation until they attain a maximum weight [53,70,72], and that the time to pupation and the actual size of the adult is affected by the temperature of the larval environment [25,61,70,72,74] as well as food level and density (see above), the type or quality of the food source [6,24,25,75–80] and, of course, the sex of the individual [21,37,70–72,74,81].

Biochemical investigations of the triggers to pupation in 4th instar larvae reveal that during the growth period of the 4th instar, more than 75% of the larval mass is accumulated [70,71,74] as well as most of the sexual dimorphism in mass [70,72]. Protein, sugars, and glycogen increase linearly with mass for both sexes [70,71,81–84] while lipids increase exponentially, and faster in males than females [70,71,81]. The triggers to pupation remain obscure, but minimum size, nutritional state, multiple hormone levels, specific gene activation/deactivation, and interactions among all of these have been implicated [25,61,70–74,81,85]. Pupal size is positively related to adult size, longevity/survival, sperm production, blood meal consumption, the size and number of eggs [1,54,73,81,84–87] and inversely related to the length of the gonotrophic cycle, and susceptibility to disease [81,87–91], but see [92–97].

Mosquito larvae in these microcosms create a dynamic system. Eggs were hatched by immersion and 1st instar larvae were counted into vials with a large number of newly-added yeast particles. As the larvae filter the particles and pass them through their guts, small initial differences in size and opportunity develop into larger differences in size. Larger individuals filter more effectively than smaller ones, so larger individuals obtain more food particles than smaller ones. However, as individuals grow, their metabolic needs increase and less of the food is used for growth as more is used to maintain the existing mass. In isolation, each individual would follow a sigmoid growth curve determined by the initial quantity of food in the vial. In the experimental vials, larvae compete for food particles with each other. There are three environmental conditions that change over time: the larvae increase in size, with concomitant increases in demand for nutrients, ability to filter, and volume of gut in which to retain particles; the number of particles remains constant, so the apparent number of particles decreases; the quality of the particles decreases with each pass through the gut of a larva. Each of these environmental conditions increases the competition among larvae as they grow. Food particles are plentiful and of high quality for the 1st instar larvae and probably for 2nd instar larvae, but become less plentiful and of lower quality for 3rd instar larvae and even less plentiful and of even lower quality for 4th instar larvae. When food particles are abundant and their quality is high, larvae pass the particles rapidly through their guts and extract the most available nutrients. During the third and fourth instars the larger size of the larvae increases their demand for food, and the relative quantity and quality of the food particles decreases. In response to this, the female larvae retain the particles in their guts for longer; this further decreases the apparent number of food particles and their quality, reinforcing the retention of food particles. The initial conditions of the experiment probably have little effect on the growth of the first and second instar larvae, but increasing effects on the third and fourth instar larvae, influencing the age and mass at pupation. Food level and density have been shown to affect competition among mosquito larvae already; the aim of this paper is to understand the differences in competition among male and female larvae and how that affects pupation and the adult life cycle.

Survival in the food x density experiment is one potential measure of competition among the mosquito larvae. In the significant interaction contrasts, Survival was highest in the vials with the least competition and lowest in the vials with the least total food, so there is an effect of competition on the survival of larvae. However, only 63 % of the variation in Survival is explained by the treatments and 30 % of that is explained by Food level alone. The MANOVA correlation coefficients corroborate the relatively low contribution of Survival to the significance of the contrasts.

The main treatments, food level and density, and the interactions, total food per vial and non-lethal competition, affect the mass and age at pupation; large mass and early pupation increase the fitness of the adult male mosquito, while large mass at pupation is primarily important to the fitness of the female mosquito [1]. Mass and age at pupation together describe the growth rate of the larva to the pupation endpoint.

### Differences between males and females

The MANOVA gives us insight into the relationships among the variables. Increased food level (mg/larva) increases mass for all larvae. Females are affected more than males, Prime females more than Average females, Average males more than Prime males. Increased food level has no effect on Prime female age at pupation and only a small effect Prime male age at pupation. Prime male age at pupation increases with increasing food level.

Also, in the MANOVA, increased density (larvae/vial) increases mass for all larvae. Males are affected more than females, Prime males more than Average males, Prime females more than Average females. Increased density increases age at pupation for both Prime males and Prime females; Prime male age at pupation is affected more than Prime female age at pupation.

There are three significant interactions in the MANOVA; these interactions describe how competition and total food per vial affect the interplay of the food level and density treatments. The three main MANOVA interactions (F1 X D3, F2 X D1, and F2 X D3) affect male mass more than female mass, Average males more than Prime males, and Prime females more than Average females. The interactions have no effect on Prime female age at pupation. Only the F2 X D1 interaction affects Prime male age at pupation.

The univariate r squared values for food level and for density are similar to the MANOVA correlations for the relationships between male mass and female mass. The mean values (mg) of all four mass variables increase with increasing food level. Density does not have the same effect on each of the four mass variables. Prime male mass is highest at the highest density. Average male mass is highest at the lowest density. Prime female mass increases with density, but is lowest at the highest density. Average female mass is also lowest at the highest density. In the univariate analyses, food level alone accounts for more than half of the variation in the two female mass variables and just less than half of the variation in the two male mass variables. Density alone accounts for much less of the variation than food level in all four mass variables. The interactions between food level and density account for more of the variation in these variables than density alone.

The r squared values for Prime male and Prime female age at pupation are relatively higher than the corresponding MANOVA correlations for both the food level and density treatments. There are effects of the experimental treatment on age at pupation that did not contribute to the MANOVA significance. The Prime male and female age at pupation are both highest at the highest food level. They are also both highest at the highest density. These are the vials with the highest total food per vial. Increased mass at pupation is beneficial to the fitness of the adult mosquito of both sexes. Increased age at pupation is potentially detrimental to adult males, probably less so to adult females. Total food per vial appears to affect the larvae and some of the effect of density may be due to the total food per vial. This should be apparent in the interactions.

The r squared values for the interactions F1 X D3 and F2 X D3, correspond to the MANOVA correlations: they explain the variance in male mass more than female mass, Average males more than Prime males, and Prime females more than Average females. The r squared values for the interaction F2 X D1 (low density treatments) are slightly higher for females than for males, and explain the same amount of variance in the mass of Prime females and Average females, and in the mass of Prime males and Average males. In contrast to the MANOVA correlations, there is a significant interaction (F1 X D3) for Prime female age at pupation, and all three interactions have significant r squared values for Prime male age at pupation.

All three of the main interactions consist of 4 treatment combinations: least competition, most competition, least food per vial and most food per vial (Tables 4-7). For instance, Survival is highest in vials with the least competition and lowest in vials with the least food per vial. Both are low density treatments. If competition is affecting the mass at pupation, the mass should be highest in the vials with the least competition. This is only true for the Average male mass and the Prime and Average female mass in the F2 X D1 interaction (low density). Alternatively, competition could be causing the lowest masses to occur in the vials with the most competition. This is the case for the Prime male and the Average male in all three interactions, and the Prime and Average females in two of the interactions (F1 X D3 and F2 X D3). Males and females respond differently to the high and low competition treatment combinations and females respond differently in low densities (D1) than across the full range of densities (D3). If total food per vial is important, then the lowest masses should be in the vials with the least total food; this is only true for Prime and Average females in the F2 X D1 interaction. Alternatively, the highest masses could be in the vials with the most total food. This is true for the Prime males in all three interactions and the Prime and Average females in the F1 X D3 and F2 X D3 interactions.

Prime males, Prime females and Average females grow largest in the vials with the most total food and smallest in the vials with the most competition. However, females at low density grow largest in vials with the least competition and smallest in vials with the least total food. Average males grow largest in the vials with the least competition and smallest in the vials with the most competition. Prime and Average females respond to the treatment conditions similarly to one another, but the Prime and Average males respond to the same treatment conditions differently from each other. Competition among males differs from competition among females. The food level and total food per vial affect competition for both sexes, but these effects are different across sexes.

Prime male age at pupation is greatest in the vials with the most food and least in the vials with the most competition. This corresponds to the Prime male mass. In the vials with the most food, the Prime male grows largest and delays pupation. In the vials with the most competition, the Prime male is smallest and pupates earliest. For the one significant interaction affecting the Prime female age at pupation, the greatest age at pupation is in the vials with the most food and the earliest is in the vials with the least competition, another difference between males and females.

Age and mass at pupation together are a measure of growth rate. For both the Prime male and the Prime female, the growth rates are highest in the vials with the least competition. This highest growth rate for the Prime female is higher than the growth rate for the Prime male in each of these interactions. Despite taking longer to pupate, females grow faster than males in the vials with least competition. [Note that growth is expected to be a sigmoid curve in which the instantaneous growth rate increases to an inflection point and then decreases towards zero. If the males pupate lower on the curve than females, then they could appear to have a lower growth rate than females even if they grow at the same rate.] The slowest growth rates for males and females are in the vials with the highest densities (most competition and most food per vial). The F2 X D1 interaction differs in that the Prime female has the lowest growth rate in the vials with the least total food.

There are consistent differences between male and female larvae across the three main interactions. Competition among males and among females needs to be considered separately. Furthermore, the results of the sex ratio experiment show that the percent of males in the vial affects the growth of males at low food levels, but not that of females; this supports the observation that there is a difference in the way the two sexes compete for the same food resource.

### Competition among females

The mass at pupation of females is directly related to the food level. Food level explains 8-10 times the experimental variance that density explains and three times the variance that the three main interactions explain (Tables 2 and 3). In contrast, age at pupation is better explained by density than by food level or the interactions, although the Prime female age at pupation is the variable least affected by the experimental conditions. Competition is described by the interactions, so competition appears to be less important to females than the food level.

These two experiments look at competition among females in five ways: the interactions identify differences in 1) mass and 2) age at pupation under various levels of competition; 3) the difference between the Prime and the Average female masses indicates possible interference competition; 4) the differences in growth rate of the Prime female in the different treatment combinations is another measure of competition; and 5) the lack of effect of the percent of males on the mass of females indicates that females outcompete males.

1) The mass at pupation responds differently in the F2 X D1 interaction than in the other two. All the vials in this interaction have either 4 larvae or 5 larvae; this is the low density treatment in the F1 X D3 and F2 X D3 interactions. Because the food is added on a per larva basis, the total food levels in these vials are lower as well. For this interaction, the largest females are in the vials with the least competition and the smallest are in the vials with the least total food (for both Prime and Average female mass at pupation). These are the vials with 4 larvae. In the higher density treatment, the vials with 5 larvae, the masses at the high food level are lower, but the masses at the low food level are higher, than in the vials with 4 larvae. The addition of a larva plus an increment of food reduces the size of the females at the high food level, but increases the size of females at the low food level. In the F1 X D3 and F2 X D3 interactions, the largest females are in the vials with the most total food and the smallest females are in those with the most competition; these are both the high density treatments (7 larvae or 8 larvae per vial). The masses of Prime and Average females in the low density treatments (4 larvae or 5 larvae per vial) are similar to the masses observed in the F2 X D1 interaction, suggesting that the difference is due to the higher density or higher total food level in the other vials. The masses of females in the F1 X D3 and F2 X D3 interactions are higher than that of the females in the F2 X D1 interaction in the vials with the greatest total food, and lower than those in the F2 X D1 interaction in the vials with the greatest competition. The additional larvae in the high density vials, plus the additional increments of food, increase the pupal masses of females in the high food vials and decrease the masses of females in the low food (most competition) vials. For females, competition is affected by food level (food/larva), density (larvae/vial) and total food per vial.

2) The Prime female pupates earliest in the vials with the least competition and latest in the vials with the most total food per vial. The Prime female age at pupation is less affected by competition (or any treatment) than any other variable.

3) There is no indication of interference competition based on the size distribution of females at pupation. At the lowest total food levels this distribution is compressed rather than elongated. In the other vials the difference between the Prime female and the Average female mass is approximately the same (0.20 mg). Comparing the Prime females in the vials with the lowest total food to those in the vials with the most competition (equivalent food/larva) and similarly comparing the Average females in those two treatments, it appears that the tighter distribution of sizes is due to the relative increase in size of the Average females at the lower density. Two things are likely causes: 1) the number of particles becomes limiting to the females earlier in the vials with the lowest total food and they switch to retention before they have developed the same size distribution as in the other vials, and 2) there is little or no difference in the ability of females to extract nutrients from the retained particles, so they all grow equally well from that point. When the females in the low total food vials switch to retention from filtering, the net effect is a more equal distribution of food among them, as compared to females in the vials with the most competition.

4) The growth rate of the Prime female is greatest in the vials with the least competition. Growth rate was approximated by dividing the mean values of the Prime female mass by the corresponding mean value of the Prime female age at pupation for each treatment combination. No additional significance tests were applied.

5) Female mass at pupation is unaffected by the percent of males in the vial (at the food levels and densities tested). Females larvae appear to be unaffected by competition with male larvae.

At low density the least competition results in the largest females and the least total food results in the smallest females. At high density the most total food results in the largest females and the most competition results in the smallest females. The low density treatments across all three interactions are comparable in treatment conditions and outcomes, so the high density (7 larvae or 8 larvae) is qualitatively different. Competition among females at high density results in smaller masses at pupation despite equivalent food/larva levels. High total food levels at high density result in larger masses at pupation despite equivalent food/larva levels. If total food per vial were entirely responsible for these observations the addition of a larva (from 4 larvae to 5 larvae) and an increment of food (per larva) should not lower the mass at pupation of females at the high food level (more total food) and increase the mass of females at the low food level (more competition). The 5 larvae in the vials with the most competition do better than the 4 larvae in the vials with the same food/larva, while the 5 larvae in the vials with the most food don’t do as well as the 4 larvae with the same food/larva, the vials with the least competition. At the higher densities (7 larvae or 8 larvae), the reverse is true. The 7 or 8 larvae in the vials with the most competition do worse than the 4 or 5 larvae in vials with the same food/larva, while 7 or 8 larvae in the vials with the most food do better that the 4 or 5 larvae in the vials with the least competition (equivalent food/larva).

For female larvae, there are 6 environmental conditions indicated by the three main interactions:

1. Vials with the least competition—These females grow fastest and pupate earliest. This is the benchmark to compare the other treatments against.
2. Vials with the least total food—These females pupate at a smaller mass than those in vials with the least competition and there is a tighter distribution of masses (smaller difference in size between the Prime and Average females) than in all the other vials. There is no indication of interference competition; the reduced size distribution is probably due to females retaining the particles in their guts for longer periods to extract more nutrients. In these vials, it appears that the low total food causes females to switch from filtering to retention at an earlier instar before large differences in size have developed, resulting in the compressed size distribution. Retention supplies fewer nutrients over time than filtering particles and passing them rapidly through the gut, resulting in smaller size at pupation and a slower growth rate than in the vials with the least competition.
3. Vials with the most total food (5 larvae)—These females have more total food than those in the vials with the least competition (4 larvae), but they don’t grow as fast or as large. The addition of one more larva even with the incremental food/larva reduces the growth rate and final mass. The larvae filter rapidly for long enough to develop the same size distribution as in the vials with the least competition, but end up approximately 0.10 mg smaller than those in the vials with the least competition. This also suggests that filtering promotes growth better than retention.
4. Vials with the most competition (5 larvae)—These females also have more total food than those in the vials with the least total food (4 larvae), and they grow larger than those females. In this case, the incremental food/larva is more beneficial than the addition of the extra larva is detrimental. Vials with 4 larvae get 8 mg or 16 mg of food, while the vials with 5 larvae get 10 mg or 20 mg of food. The mean Prime and Average female masses for total food levels of 8 mg and 10 mg range from 2.50 mg to 2.52 mg. The masses for total food levels of 16 mg and 20 mg are 1.0 mg higher (40%, 3.45 mg to 3.80 mg). The largest increase is in the vials with 5 larvae and 20 mg total food. There appears to be a change in the growth of female larvae at food levels between 16 mg and 20 mg total food per vial that results in a disproportionate increase in the mass of females. It appears that females in vials with less food switch to retention as they perceive the number of particles decreasing and thereafter grow more slowly, while females in vials with more total food (20 mg per vial and greater) continue to filter and grow at a faster rate, and to a larger size.
5. Vials with the most total food (7 larvae or 8 larvae)—These females grow larger than any other females and take longer to pupate. All these vials have more than 20 mg of food in them. Females in the vials with the least competition grow faster and pupate earlier, so these females delay pupation and become larger. As mentioned earlier, larger filter feeders have an advantage over smaller ones, but as they grow in mass, each increment in mass adds less and less to that advantage. At some point the individual will reach an equilibrium where the filtering is only sufficient to maintain its mass, not increase it. Females in vials with the most total food may grow as fast as those in the vials with the least competition and then extend their larval growth period to increase in size at the expense of their growth rate. For females, large adult mass is more beneficial than early age at pupation.
6. Vials with the most competition (7 larvae or 8 larvae)—These females are smaller than those in any other vials. The total food in these vials ranges from 14 mg to 32 mg (14 mg, 16 mg, 21 mg, 24 mg, 28 mg, 32 mg), yet the females pupate at smaller masses than those with the least total food: 8 mg to 20 mg (8 mg, 10 mg, 12 mg, 15 mg, 16 mg, 20 mg). The total food per vial is higher and the relative food (per larva) is identical, yet the females in the higher density vials do not grow as large. These females must switch from filtering to retaining particles later than those in the vials with the least total food, because they do develop a size distribution that resembles the vials with the least competition and the vials with the most total food. The additional larvae in the vials with the most competition must reduce the available particles sufficiently that the effective food level is lower than in the vials with the lowest total food.

#### Summary

The environment that the larvae experience in their vials changes over time and the larvae respond depending on the initial conditions of the vials (food level and density). For females, food level (mg/larva) is the most important factor, but competition, total food per vial, and density also influence the mass and age at pupation. Females in the vials with the least competition grow fastest. They pupate at large sizes (not always the largest, but close) and earlier than in the other treatments. These vials are the optimum environment for females within this experiment. These females probably filter particles and pass them rapidly through their guts until they pupate. Pupation probably occurs when the quality of the food particles is insufficient to support further growth. In contrast, females in the vials with the least total food begin retaining food early in larval development. They grow slowly and are among the smallest at pupation. Pupation probably occurs when the quality of the food particles is insufficient to support further growth. Females in vials with 5 larvae (instead of 4 larvae) don’t grow as large at the higher food levels, so the added larva reduces the apparent number of particles and causes a switch to retention before pupation. Retention is less effective than filtering so these females are smaller than those in the vials with the least competition despite the equivalent food/larva. However, females in vials with 5 larvae (instead of 4 larvae) grow larger in the vials with the most competition. In this case the total food per vial is greater and the females switch to retention later than those in the vials with the least total food. They don’t grow as large as the females in the vials with more total food, but they are larger than the ones with the least total food (despite equivalent food/larva). At higher densities (7 or 8 larvae) the females in the vials with the most competition are even smaller at pupation than those in vials with the least total food. These females switch to retention later than those in the vials with the least total food; they develop a distribution of sizes similar to those in the vials with the least competition. Once they switch to retention, the larger number of females reduces the apparent number of particles below the level of that in the vials with the least total food, and the pupae are smaller despite the equivalent food/larva. In these vials, pupation may be triggered by low food particle quantity rather than low quality. Again at high densities (7 or 8 larvae) the females in the vials with the most total food are even larger than those with the least competition. These females filter and pass the food rapidly through their guts until pupation, similarly to those females in the vials with the least competition. However, the large excess of food allows them to continue to grow, albeit at a slower pace, for more than a day after the Prime females in the corresponding vials with least competition have pupated. It is possible that pupation is triggered by larval size rather than by diminished food quality or quantity.

### Competition among males

Male larvae respond differently to the treatment conditions and interactions than female larvae. While male pupal mass is also affected by the food level more than by density or interactions, density and interactions are relatively more important than food level as compared to females. Food level explains only 41 % to 47 % of the variance in mass at pupation for males, compared to 64% for females. Density explains 12 % to 16 % of the variance in mass at pupation for males compared to 6 % to 8 % for females. The three main interactions explain 27 % to 31 % of the variance for males compared to 20 % to 22 % for females (Tables 2 and 3). These comparisons are qualitatively similar to the MANOVA results in Table 1. Since competition effects show up in the interactions, competition is relatively more significant for males than for females, and more significant for the Average males than for the Prime male. Another difference between males and females is in the way that the Prime male and Average males respond to the interactions. While both Prime and Average females’ masses respond to the interaction treatment combinations in the same way, the Prime male and the Average male masses grow largest in different treatments. The Prime male mass is largest in the treatments with the most total food per vial, similar to the female mass variables, while the Average male mass is largest in the vials with the least competition.

These two experiments look at competition among males in 5 ways: the interactions identify differences in 1) mass and 2) age at pupation under various levels of competition; 3) the difference between the Prime and the Average males indicates possible interference competition; 4) the differences in growth rate of the Prime male in the different treatment combinations is another measure of competition; and 5) the significant regression of the percent of males on the mass of the Prime males at the lower food level.

1 & 2) The Prime male grows to the largest mass and takes the longest to pupate in the vials with the most total food. The Prime male pupates earliest and at the smallest mass in the vials with the most competition. These are both high density treatments. The vials with the least competition produce Prime males that are smaller than the ones with the most total food, and they pupate earlier, however, Prime males in the vials with the least competition have the highest growth rates across all the vials. Prime males in all three interactions grow and pupate like Prime and Average females in the F1 X D3 and F2 X D3 interactions; total food per vial has a positive effect on mass at pupation and competition has a negative effect. However, the effect of the treatments on the age at pupation is different for males and for females; males delay pupation at high total food per vial, like females, but pupate earliest in vials with the most competition, where females pupate earliest in vials with the least competition.

Average males grow largest in the vials with the least competition and are smallest in the vials with the most competition. Another feature of the vials with the least competition is that the Average male mass is greater than the Prime male mass; the male larvae remaining in the vial after the Prime male pupates grow larger than the Prime male on the food resource that is left. Considering that the Prime male outcompetes the other males, when the Prime male pupates, the next larger male should assume the dominant role until he pupates, followed by the next, and so on. At least one of these males must be larger than the Prime male for the Average mass to be greater than the Prime mass. This implies a competitive release after the Prime male pupates as well as a considerable amount of food left over. Males in the vials with the highest total food per vial take longer and grow larger than the males in the vials with least competition. If optimizing mass against early pupation were the only criteria that male larvae use to determine when to pupate, then the Prime male mass and age at pupation should be the same here as in the vials with the least competition. They are not; males in a high food environment delay pupation and increase further in mass.

3) There is no indication of interference competition based on the size distribution of males at pupation. At the lowest total food per vial this distribution is compressed rather than elongated as it would be if the larval mosquitoes engaged in some kind of interference competition. In vials with higher densities the difference between the Prime male and the Average male mass is approximately the same regardless of the treatment. This smallest difference in size between the Prime male mass and the Average male mass in the vials with the lowest total food per vial may indicate an effect of total food per vial at the lower end of the range as well as at the upper end. The Prime male in these vials is not much larger than those in the vials with the most competition and the age at pupation is also not much greater. However, the Average male mass in these vials is relatively larger than the corresponding masses in the vials with the most competition, resulting in the tighter size distribution. More intense competition for food due to the particle retention by the female larvae reduces the size of the Prime male, but allows the non-Prime males to continue to grow after the Prime male pupates, whereas the most intense competition affects the non-Prime males more.

4) Prime males in the vials with the least competition have the highest growth rates across all the vials. Prime males in the vials with the most competition take less time, and pupate at lower masses than those in the vials with the least competition. They also pupate earlier and at lower masses than Prime males in the vials with the least total food per vial (the same food/larva treatments at the lower density). The effect of increased density is to increase competition despite increasing the total food per vial and maintaining proportional food resources. Under this competitive stress Prime males pupate earlier and at lower masses. The growth rates are also lower, suggesting that the males are smaller not just because they pupate earlier, but because they are not able to grow as well.

5) From the sex ratio experiment we know that at low food levels, the Prime male mass increases as the percent of males in the vial increases. The Average mass does not change so the increase in mass of the Prime male is offset by the decrease in mass of the non-Prime males. The Prime male outcompetes the other males for food (at 3 mg/larva) and the advantage of the Prime male increases as the percent of males increases.

Because the Prime male benefits from the increased percent of males at the low food level, but none of the other mass variables are affected by the percent of males at either food level, the environment that males experience in the vials must take into consideration what the female larvae are doing. During the early growth of the larvae, food levels are expected to appear high because the larvae are small and the quality of the food is initially at its highest. As they grow and females switch from filtering to retention in some vials, the males will experience a reduction in the number of food particles as well. Since the Prime male is affected by the percent of males in exactly the vials where the females are retaining food, it appears that males do not retain food but continue to filter particles and pass them rapidly through their guts even as the particle numbers decrease.

There are only 4 environmental conditions indicated by the three main interactions for males:

1) Vials with the least competition—Prime males grow at the fastest rate and pupate at sizes close to the largest. Average males grow even larger. Since females in these vials also grow at the fastest rate across the experiment, it appears that the Prime male filters particles and grows to a size that allows or triggers pupation, and then the remaining males experience a net increase in food particles that allows them to grow even larger. The Prime male is not retaining particles as females do when particle numbers or quality decreases, but it does sequester some number of particles as they pass through the gut. It is the release of these particles at pupation that drives the non-Prime males to grow further.

2) Vials with the least total food—the Prime male pupates at masses and ages that are almost as low as in the vials with the most competition. In these vials, the females appear to be food limited and switch from filtering to retention earlier than in other vials, reducing the number of particles and the particle quality further. These vials correspond to the conditions in the second experiment where the Prime males benefit from the increased percent of males in the vial. The Prime male competes with the other males for particles, the numbers of which the females affect by retaining the particles in their guts. The low total food per vial causes the females to switch to retention earlier and this causes the males to experience an even lower total food per vial. The Prime male pupates at a size almost as small as in the vials with the most competition and almost as early. As in the vials with the least competition, the non-Prime males experience a small benefit from the additional food made available once the Prime male pupates, and they grow larger than the non-Prime males in the vials with the most competition. While the size distribution of the female larvae is compressed because they switch to a less effective method of feeding at low food levels, the size distribution of the male larvae is compressed because they have less total food available to them due to the retention of the females. The results of the sex ratio experiment indicate that the males are competing in a pure exploitative mode at the low food levels (where one would expect interference competition), the fewer the females present, the more available particles and the larger the Prime male grows. Because the size of the Average males is not affected by the percent of males, the non-Prime males decrease in size proportionately to the Prime male’s increase. This indicates that the males are filtering even at low food levels. If they were retaining food particles as the females appear to do, we would not see this change in the size distribution due to the percent of males in the vials.

3) Vials with the most total food—in these vials, females extend their growth beyond that of females in the vials with the least competition and Prime males do the same thing (both age and mass at pupation are greater). Prime males grow larger and pupate later in these vials than in any others across the experiment. The females filter particles throughout their larval growth and extend that growth period for two days longer than the Prime male. Prime males pupate between 5 and 6 days except in these vials with the most total food. Prime females pupate between 6 and 9 days except in the vials with the highest densities. When food particles are available and/or food quality is still high, the 4th instar larvae of both sexes delay pupation to increase further in size. The Prime male is 0.5 mg to 1.0 mg larger than Prime males in the vials with the least competition.

The Average male in the vials with the most total food are almost as large as the Average males in the vials with the least competition. They are similar in size to the Prime males in the vials with the least competition (.03 mg larger to .05 mg smaller) indicating that males experience little competition in the vials with the most total food. However, they do not experience the release of food particles and grow to be larger than the Prime male as in the vials with the least competition. The most likely explanation for the difference in outcome between the Average males in the vials with the most food and the Average males in the vials with the least competition is that males are constrained or driven to pupate by a certain age so when the Prime male delays pupation it compresses the distribution of ages at pupation for the rest of the males, limiting the benefit that the additional food bestows on the non-Prime males in the vials with the most total food. [The distribution of ages at pupation within microcosms was not analyzed in this experiment.]

4) Vials with the most competition—the Prime male pupates at the earliest age and smallest mass. The Average male mass is also smallest in these vials. Prime and Average female masses are also smallest at the highest Density (7 or 8 larvae/vial) and close to the smallest in the 5 larvae/ vial treatment. The total food per vial is high enough at these high densities so that the larvae filter and grow large enough to develop a size distribution similar to those in the vials with the least competition. As they grow food particles become relatively scarcer and the females switch from filtering to retaining the particles, making them even scarcer. The males respond to the change in the number of food particles by pupating earlier and at the smallest sizes across the experiment.

Summary: The environment that the larvae experience in their vials changes over time and the larvae respond depending on the initial conditions of the vials (food level and density). For males, food level (mg/larva) is also the most important factor, but density, total food per vial and competition (interactions) are relatively more important than for females. Male larvae are also affected by the number and behavior of female larvae. Male larvae filter particles and pass them through their guts, extracting the most available nutrients; they do not appear to change this feeding strategy to retain particles at the expense of their growth rate, as female larvae do. Males in the vials with the least competition grow fastest, and pupate at large sizes, especially the nonPrime males, which pupate at masses larger than the Prime male in each vial. This indicates a release from competition for the non-Prime males when the Prime male pupates. Males in the vials with the least total food experience a reduction in food particles when the females begin retaining food and develop a similar compressed size distribution to those females. They pupate relatively early and at a relatively small size (not the smallest, but close). Males in the vials with the most competition pupate earlier and at smaller sizes than in any other vials; they appear to have less food available to them than those in the vials with the least total food, despite equivalent initial food/larva. The quality and quantity of particles is reduced by the behavior of the female larvae and this causes the males to pupate early and at a small size. In the vials with the most total food, Prime males grow to their largest sizes and delay pupation by a day to attain that size. Average males grow large as well, but not as large as they do in the vials with the least competition. This is possibly because of the delay in pupation by the Prime male; there may be a time constraint on pupation that reduces the benefit from the release of competition observed in the vials with the least competition.

### Competition between males and females

Female mosquito larvae grow to be larger than male larvae in similar larval environments. Across the experiment, some Prime and Average males are larger than some Prime and Average females, but within any vial, the Prime and Average female masses at pupation are always larger than the corresponding Prime and Average male masses (Tables 4C, 4E, 4F, 4H). Comparing the pupal masses across treatment combinations in the interactions shows a larger difference in size between females and males (Prime vs Prime, Average vs Average) in the high food/larva treatments than in the low food/larva treatments. In the F2 X D1 interaction the difference in size parallels the results of the female mass: the largest difference is in the vials with the least competition and the smallest difference is in the vials with the least total food. These treatments are both at the low Density (4 larvae/vial). The vials with 5 larvae show intermediate differences between the females and males, with the higher Food level also having the greater size difference. In the F1 X D3 and F2 X D3 interactions the greatest differences in size between the females and males are in the high Density (7 or 8 larvae) vials with the most total food, and the smallest differences are in the high Density (7 or 8 larvae) vials with the most competition. All these differences reflect the mass of females; in vials where the females grow largest, the difference between males and females is largest, and in vials where the females are the smallest, the difference between males and females is also smallest. This shows up in the MANOVA and the ANOVA r squared results (females are affected by Food level more than males). Females vary in size according to Food level and total food per vial with competition limiting pupal mass at high densities. Males vary less in size than females and they are smaller and pupate earlier. Males also vary in size according to Food level and total food per vial, but Density and interactions account for 50 % or more of the variance, and competition is important at both low densities and high densities, at least for the non-Prime males. Within the limits of these experiments, females control the availability of food; males, especially the Prime male, escape competition by growing as rapidly as possible and pupating as the available food decreases.

In the F1 X D3 interaction, the difference between Prime females and Prime males in the vials with the least competition is as great as that difference in the vials with the most food. Prime males and females grow at the fastest rate in the vials with the least competition; these vials are expected to be the optimal conditions for the larvae within this experiment. The large difference in size between the Prime female and the Prime male in these vials is the result of optimal growth. Both Prime females and Prime males grow larger in the vials with the most food per vial, but the difference between them remains the same. This suggests that even at very high total Food levels there is an optimal largest size for both males and females. Furthermore, the incremental size between Prime females in the vials with the least competition and those in the vials with the most food is only 0.08 mg, while the incremental size between males in the corresponding vials is only 0.10 mg. Prime females delay pupation for 1.53 days to achieve this incremental growth and Prime males delay it for 0.73 days.

Again, in the F1 X D3 interaction, the difference between the Prime females and Prime males in the vials with the least food per vial is as small as that difference in the vials with the most competition. From the second experiment we know that the Prime male benefits from higher percent males under the conditions in the vials with the least food per vial and that the non-Prime males are smaller. The differences between the Prime females and Prime males in these two treatments are smaller than in any other vials across all three interactions. The sizes of the Prime female and Prime male in the vials with the most competition is even lower than their sizes in the vials with the least total food. This suggests that competition in these vials has a greater deleterious effect on growth than a lower total food per vial, which compresses the size distribution of both males and females. In the vials with the most competition, the size distribution is comparable to the other vials at high Density; this means that the non-Prime males and non-Prime females are reduced in size by competition more than they are in the vials with the least total food.

If there were interference competition between males and females, the size distributions should be larger at lower Food levels rather than smaller as observed. Competition between males and females is exploitative, but females control the food resource in two ways: female larvae grow larger than males and dominate the competition for particles by filtering faster; and female larvae retain food particles when food becomes relatively scarce so that the available particles become even scarcer. The Prime male pupates within 5 to 6 days after hatching except when food availability is high and it extends larval growth to increase in size at the expense of its growth rate. The Average male grows to a size dictated by the initial Food level and competition. The Prime female pupates within 6 to 8 days after hatching except when Density is high and it extends its larvae growth. Two of these exceptions are at the highest Density and lowest Food levels (most competition) and two of them are at the highest densities and highest Food level (most total food), so there are potentially different causes for the extension in larval growth among females. The Average female mass at pupation reflects the growth patterns of the Prime female.

Mosquito larvae do not grow in a smooth curve; each of the 4 instars grows within the constraints of a larval exoskeleton, which is shed at the subsequent molt (see [58,63]). The two observed size distributions (compressed in the vials with the least total food, larger size distributions in the other vials) suggest that the female larvae in the vials with the least total food switch to retention in the third instar or earlier, while the rest of the vials do not experience the lowered food particle levels that trigger retention until sometime in the 4th instar. Most of the competition will then occur in the 4th instar for both males and females. Mosquito larvae do not display obvious secondary sex characteristics, but there is a bimodal size dimorphism among older (4th instar) larvae and larger larvae are usually female. By the beginning of the 4th instar at least, female mosquito larvae already have developed a competitive advantage over the males [70,72].

### Interference competition

The distribution of sizes among males and among females is another measure of competition (besides the absolute size). In tadpoles, low size and a large difference between the Prime and Average tadpoles indicated that interference competition mechanisms replaced exploitative competition at low food levels [30,31,98,99]. Rubenstein’s [100] data suggest similar interference competition among Pygmy Sunfish in the lab experiments, but not in the field trials.

There is no such pattern comparing Prime male to Average male mass, Prime female to Average female mass, Prime female to Prime male mass, or Average female to Average male mass across the three main interactions. The differences between Prime male and Average male, and Prime female and Average female are smallest in the vials with the least food per vial; interference competition would produce a larger difference between the Prime and Average individuals in these vials. Differences between sexes are also lower at the lower food levels rather than higher, so competition appears to be purely exploitative. However, there are differences in the patterns for males and females, suggesting that they are competing differently for the food resource.

Interference competition among *A. aegypti* larvae has been postulated based on “Growth Retardant Factor (GRF)” in conditioned water[16,41,43,45,56,101], but see [47]. That water conditioned by rearing mosquito larvae in it negatively affects the growth and/or survival of subsequent larvae is insufficient to conclude that there is interference competition among the larvae; evidence of reduced size of the original competitors and a larger difference between the Prime and Average pupae of each species is necessary as well. I did not observe interference competition in my experiments, but it is possible that the strain of *Aedes aegypti* that I collected did not produce the GRF observed in other experiments (see [47,101]).

### The effects of food and density

Mosquito larval growth determines the size of the adult mosquito. Microcosm experiments allow the manipulation of external factors that influence larval growth and reveal the effects and interactions among those factors. In these experiments the factors were food level (mg/larva), density (larvae/vial), and sex ratio (% males/vial), and the interactions showed the contributions of competition and total food/vial. There were clear differences between the responses of male and female larvae to the initial conditions of food level and density and to competition and the total food/vial. Females dominate competition for food particles in these experiments, but the responses of both males and females were more complex than prior investigations predicted.

Investigations of the larval ecology of *A. aegypti* have held the total food level constant and varied the density of larvae [4,7,10,33,36,47,63,75,90,96,102-107]. Others varied food level at a constant density of larvae [3,24,25,35,38-40,55,62,68,76,82,92,108] or varied volume and surface area while keeping the number of individuals and food level constant [40,82]. Greenough et al. [32] varied the number of individuals while keeping total food and volume proportional to the number of individuals. Serpa et al. [109] varied number of individuals but decreased both food and volume as the number of individuals increased. Mitchell-Foster et al. [110] and Price et al. [87] also increased density and decreased food at the same time. While none of these investigations contradicts the results of my experiments, none of them can be used as direct support because their designs do not allow the possibility of an interaction between food and density.

Other investigators varied both food level and density [2,15,17-22,26,27,34,79]; these data show that food level interacts with larval density affecting the outcome of larval growth. Some of these prior studies demonstrate differences between males and females [2,15,17,20,21,32,33,79], but only Wada [15], Daugherty et al. [2], Agnew et al. [20], Bedhomme et al. [21], Kim & Muturi [79], and my own results indicate that the effect of the interaction between food level and density differs for males and females.

*Aedes aegypti* larvae occur in nature in low numbers spread across multiple small containers [2-13,111], but see [14]. There are likely to be only a small number of winners in each of these competitive arenas and the Prime male and Prime female are proposed as good proxies for these ecological and evolutionary winners. Average values of pupal size and age at pupation are not the best estimates of the outcome of competition in the containers. The mosquito larvae react to their environmental conditions including: food level (food/larva), total food (food/container), density (larvae/container), and competitive interactions differently depending on gender as well as these conditions. The outcomes form a complex pattern over the conditions tested, which were preselected for non-lethal competition. Based on the complexity of the observed responses to the environmental conditions, it would be unwise to infer that these results predict the outcomes beyond the experimental design; these mosquito larvae appear to behave in a complex and reproducible way despite the simplicity of the experimental treatments.

Other investigators have identified additional environmental conditions that affect the growth and survival of *A. aegypti* as larvae: temperature, temperature fluctuations, other food sources, pesticides, pollutants, parasites, predators, and competitors (see above). Furthermore, adult behavior and the ability of *A. aegypti* eggs to remain quiescent despite repeated inundations [7,10,112–119] complicate the natural history of this species and suggest that its resilience, invasiveness and association with human habitation may be due to multiple feedback cycles that allow a small, diffuse population to rebound repeatedly after eradication attempts.

Studies on other mosquitoes, especially those of interspecific competition among larvae of similar species, suggest that there may be significant differences in the ecology, physiology, and behavior of even congeneric mosquito larvae as compared to *Aedes aegypti.* (Aedes albopictus — [26,34,63,67,68,75,79,89,93,94,104,106,111,112,120-145]; Aedes sierrensis—[146–153]; Aedes triseriatus—[154-179].

### Implications for vector control

*Aedes aegypti* larvae respond to the amount of food and larval density in their containers such that low densities of larvae will produce larger pupae of both sexes compared to higher densities at any given food level (mg/larva). This suggests that vector control efforts to reduce the adult population may result in lower numbers of larvae in subsequent generations and thus larger pupae and adults.

Larger adults may be more robust and longer lived than smaller ones and larger females may take larger blood meals, produce more and larger eggs, [1,81,92] and possibly survive to take a second blood meal [91]. Each subsequent blood meal is an opportunity for the transmission of Zika, Dengue, Yellow Fever or other diseases transmissible by A. aegypti. Adult females bite multiple times per blood meal in the laboratory (personal observations), so a female may be able to transmit a virus from one individual to another even during the initial blood feeding cycle. There may be a difference in risk to human health between a large population of small mosquitoes versus a smaller population of larger mosquitoes, but neither option is desirable. The best outcome of control efforts may be to remove larval habitat rather than to attempt to control adult populations.

### Implications for mosquito ecology

Many investigators have asserted that *Aedes aegypti* larvae are food limited in their normal environment and some investigators have observed this to be true [3,37,180–182]. Other investigators have observed that *A. aegypti* larvae are not always food limited in their normal environment [4,7,10,183,184].

*Aedes aegypti* exhibits a number of ecological adaptations that make it resilient in the face of an uncertain environment. In addition to the competitive responses of the larvae to food and density, which consistently produce early maturing males and larger, later females, the larvae can survive without food for long periods waiting for additional food input so as to complete four instars and pupate (almost 25 days for a 4th instar larvae at 20 C, less for earlier instars and higher temperatures, [74]; see also [18,75,183]. Adults can survive on sucrose, a nectar substitute, in the lab for 80-105 days [86,108,138,185]. Adult females require a blood meal to mature eggs. The size of the adult female is directly related to the size of the pupa and reflects the competitive success of the larva; large females take larger blood meals, mature more and larger eggs, and live longer than small females [1,4,18,23,35,58,70,81,83,84,86,104,110,129,186,187], but see [35], and may be more effective disease vectors [88–91], but see [92–97,188] although this is likely a result of their robustness rather than a direct result of size selection. Larger eggs hatch into larger larvae and start their lives with an advantage [54]. Large females may also fly further than small females, so may have more oviposition sites (containers) available to them [86,187], but see [35]. Recaptured adult females fly as far as 200 m [35] from their release point (but see [86] for lab flight potential >1 km); longer distance dispersal may be primarily as eggs in containers transported by humans [118,189]. However, small females may also mature large eggs, giving their offspring an advantage similar to that of large females [54,81]. Females deposit their eggs individually and may use multiple containers; the criteria that female *A. aegypti* use to select the containers and to allocate the number of eggs across containers includes presence of conspecific larvae and pupae, container fill method, container size, lid, and sun exposure [8,60,113] and the presence of other species’ larvae and pupae [118]. Another factor in the resilience of this species is the ability of eggs to survive for some time until they are inundated, and the additional feature that not all of the eggs hatch during the first inundation, forming a reserve egg pool in the event that the initial hatch is unsuccessful [7,10,112–119].

Adult male *A. aegypti* emerge before their sisters. This may reduce inbreeding in this species composed of many small localized populations (spatially and temporally). Size seems to be less important to males than to females [1,54], but larger males produce more and better sperm (and seminal fluid proteins) and live longer than smaller males [35,185,190], but see [191]. There doesn’t seem to be size selection at mating for either males or females [1] suggesting that low population densities and short adult lives constrain the opportunities for mating and offset any advantage that size selection might confer. Despite this, there is a clear positive feedback effect reinforcing the value of large size at every life cycle stage.

*Aedes aegypti* can survive for relatively long periods as eggs, starving larvae, or sugar feeding adults while waiting for an opportunity to hatch, feed, mate, and blood feed, so as to progress to the next stage of the life cycle. Large size confers an advantage to eggs, larvae and adults, but since size is a plastic response to larval conditions, it doesn’t seem to affect mating preference (but see [37] on the heritability of size). Timmermann & Briegel [71], and Price et al. [87] suggest that two mosquitoes of the same size with different larval (nutritional) histories may not be equal, with differences in their internal reserves and metabolic capacities. The physiology of nutrient accumulation and its relationship to the hormonal triggering of pupation have been studied, but the results seem contradictory (see above). The ecological evidence and the results of the current study suggest that these physiological triggers (responses to environmental conditions) might be much more complex than currently understood. In the current study, the initiation of pupation appeared to be affected by food level (quantity and quality of particles), larval density, total food/vial and competitive interactions with other larvae, as well as by larval mass, nutritional history (inferred) and sex. Larval competition appears to be more important to determining pupal size and timing for males, while food level and total food/vial appear to be more important to females. Furthermore, these environmental factors may change the triggers for pupation in different ways for each of the sexes. Two other factors that may influence pupation are the distribution of sizes within each sex (the difference between the Prime and nonPrime individuals), and the absolute size for large females (i.e. there may be a maximum size at any set of ecological conditions: temperature, food type, larval density, etc.).

### Evolution

This species has been spread globally by inadvertent human activity and has been actively eradicated for decades. Nevertheless, it persists in small, relatively isolated, and impermanent populations using various water-filled containers as a larval resource and vertebrate hosts as an adult protein resource to produce eggs. Differences in strains have been observed [18,101,192], but de Lourdes Munoz et al. [193] found that proximity did not result in genetic similarity along an 800 km range. The evolutionary implication is that this species retains its identity globally despite expected allopatric pressure to diverge because of continued transport and reintroduction by human activity.

## Acknowledgements

I would like to thank Dr. J. H. Frank for his advice and encouragement. Drs. W. Bradshaw, J. R. Linley, L. P. Lounibos, G. F. O’Meara, J. M. Werts and Mr. J. G. White also read the manuscript and offered helpful suggestions. The research was carried out while I was supported by an NIHNIAID post-doctoral fellowship (AI-06088) at the Florida Medical Entomology Laboratory. I am grateful to Dr. Werts and the Department of Zoology at Duke University for computer time. My late partner, Richard O. Merritt, provided financial support to allow me to complete the paper after many years delay. My present partner, Dr. Douglas Kline, provided encouragement, editing and technical guidance in the completion of the task.

## Supporting information

S1 Table. Sample size (number of replicates analyzed) by treatment.

S2 Table. Significant correlations between composite scores and variables with MANOVA significance levels and R squared by contrast.

S3 Table. Mean squares, significance levels, and r squared values by single DF contrast for each of the 7 dependent variables.

S4 Table. Survival (Arcsin transformation of percent survival) by treatment.

S5 Table. Prime female mass at pupation (mg) by treatment.

S6 Table. Average female mass at pupation (mg) by treatment.

S7 Table. Prime male mass at pupation (mg) by treatment.

S8 Table. Average male mass at pupation (mg) by treatment.

S9 Table. Prime male age at pupation (days) by treatment.

S10 Table. Prime female age at pupation (days) by treatment.

